# Feasibility Study of an Optical Caustic Plasmonic Light Scattering Sensor for Human Serum Anti-Dengue Protein E Antibody Detection

**DOI:** 10.1101/080218

**Authors:** A.A. Garcia, L.S. Franco, M.A. Pirez-Gomez, J.L. Pech-Pacheco, J.F. Mendez-Galvan, C. Machain-Williams, L. Talavera-Aguilar, J.H. Espinosa-Carrillo, M.M. Duarte-Villaseñor, Ch.J. Be Ortiz, L.E. Espinosa de los Monteros, A. Castillo-Pacheco, J.E. García-Rejon, B Chavez

**Author notes:** Equal contribution (A.A. Garcia, L.S. Franco).

## Abstract

Antibody detection and accurate diagnosis of tropical diseases is essential to help prevent the spread of disease. However, most detection methods lack cost-effectiveness and field-portability, which are essential features for achieving diagnosis in a timely manner in developing countries. To address this problem, transparent 3D printed sample chambers with a total volume of 700 microliters and an oblate spheroid shape were fabricated to measure green light scattering of gold nanoparticles using an optical caustic focus to detect antibodies. Scattering signals from 90 degree scattering of 20, 40, 50, 60, 80, 100, and 200 nm gold nanoparticles using a green laser and standard quartz cuvette were compared to the scattering signals from a green LED light source with an oblate spheroid sample chamber and to Mie theory by fitting the data to a logistic curve. The change in signal from 60 nm to 120 nm decreased in the order of Mie Theory > Optical Caustic scattering > standard laser 90 degree scattering. These results suggested that conjugating 60 nm gold nanoparticles with Dengue Protein E and using an optical caustic system to detect plasmonic light scattering would result in a sensitive test for detecting human antibodies against Dengue Protein E in serum. To explore this possibility, we studied the light scattering response of protein E conjugated gold nanoparticles exposed to different concentrations of anti-protein E antibody, and posteriorly via a feasibility study consisting of 10 human serum samples using a modified dot blot protocol and a handheld optical caustic-based sensor device. The overall agreement between the benchtop light scattering and dot blot results and the handheld optical caustic sensor suggest that the new sensor concept shows promise to detect gold nanoparticle aggregation caused by the presence of the antibody using a homogeneous assay. Further testing and protocol optimization is needed in order to draw conclusions on the positive predictive and negative predictive values for this new testing system.

## Background

The development of a rapid and accurate means for identifying the cause of a systemic infection has been a subject of intense interest due to the need for proper course of treatment in order to improve patient outcomes. Also, there is a growing need for accurate diagnoses due to the toll in human and economic terms caused by endemic diseases transmitted by mosquitoes in tropical regions. Of particular focus in developing countries in the tropical region, is the ability to ascertain whether a patient has contracted Dengue Fever and specifically which serotype (DEN-1, 2, 3, 4) is the probable cause of the illness. While symptomatic diagnostics play an important role in all febrile disease management, some patients are asymptomatic or have mild symptoms, and diagnosis of these patients can be critical because Dengue Fever serotypes are believed to be directly linked to severe courses of infection such as Dengue Shock syndrome (DSS) and Dengue Hemorrhoragic Fever (DHF), making early serotype identification a potentially valuable tool. Moreover, confirmation of whether the patient has had a prior serotype infection, which has also been linked to DSS and DHF, can also help to determine the course of treatment and further steps to prevent an outbreak of Dengue [8–11].

In order to meet the need for a rapid diagnostic that can be implemented in the field and closely based on patient exposure and/or immune response at an early stage of patient reported symptoms, a hybrid approach that combines gold nanoparticle conjugates (similar to what is used in low-cost paper diagnostics based on lateral flow immunoassay technology) with a high level of sensitivity in conjunction with a portable sensor for detecting a positive response at low antigen levels is the subject of this feasibility study. In order to establish a relatively low cost diagnostic tool, the hybrid technology described in this paper builds upon the employment of an optical caustic light scattering technology that employs no lenses and does not need filters [1]. Moreover, the use of battery powered LEDs and the deployment of a smartphone to analyze the scattered light are also helpful in reducing cost as well as in improving access to the device for low and middle income countries.

To commence a process of verifying the capabilities of the new hybrid technology platform in a simulated low resource setting, our approach was to first determine if an existing Dengue molecular reagent used for gold nanoparticle conjugation in paper assays could detect Dengue antibodies with human patient samples by employing this new hybrid technology platform. For the first part of this study, we analyzed the 90 degree light scattering response of conjugated gold nanoparticles to different concentrations of anti-Protein E antibody to determine key parameters such as: (1) size of gold nanoparticle that should be used; and (2) stability of the gold nanoparticle conjugates when subjected to transportation and varying temperature conditions. For the second part of the study, a direct approach with clinically relevant samples was used since it is a faster way to determine other important parameters such as: (1) whether very small amounts of human serum samples can yield reasonable signals; (2) determine the need for additional dilution or blocking steps when working with human serum; and (3) whether room or body temperature and incubation time generate differences in measurement. Thus, the overall objective of this feasibility study is solely to gain experience in practical aspects of developing a rapid and hybrid quantitative assay for determining Dengue infection by monitoring patient antibody response while simulating a low resource setting in order to expand testing in a variety of locales and with a wider variation in infrastructure than is normally the case in high income country clinics and hospitals.

## Materials and Methods

### Gold Nanoparticle Light Scattering Calibration

For calibration purposes, unconjugated gold nanoparticles of 20, 40, 50, 60, 80, 100, and 200 nm (Ted Pella Inc., Reading, CA) were diluted 1:10 in distilled deionized water and analyzed for light scattering and extinction using three separate detection methods. Each gold particle size sample contained the same overall molar concentration of gold.

### Gold Nanoparticle Conjugation

In a 1.5 ml Eppendorf tube, an aliquot of 1 ml containing 2.6 × 10^10^ particles/ml of 60 nm gold colloids (Ted Pella Inc, Redding, CA) was combined with 100 microliters of freshly prepared pH = 8.5 borate buffer prior to the addition of 50 microliters of 0.85 mg/ml Protein E Dengue Envelope-2 32 kDA (MCR-054, Reagent Proteins, Pfenex Inc., San Diego, CA). A frozen cold pack was placed on the tube and the mixture was gently rocked using an orbital table for 30 minutes. This was followed by the addition of 2.5 microliters of 10% Tween 20 and rocking on the orbital table for 5 more minutes.

One or three centrifugation steps were then conducted in order to reduce the concentration of unbound Protein E form the suspension. A Beckman Coulter Microfuge 18 (Beckman Coulter, Brea, CA) was placed in a 4˚C environmental room in order to minimize aggregation of gold nanoparticles and ensure resuspension after centrifugation. The first centrifugation was for 30 minutes at 3500g. Immediately after centrifugation, as much of the supernatant as possible was carefully removed from the pellet using a 200 microliter micropipette, while not disturbing the pellet. Once this was accomplished, 1 ml of pH = 8.5 borate buffer was added to re-suspend the pellet. An aliquot of 2.5 microliters of 10% Tween 20 was then added to the suspension and the mixture was gently rocked for 5 minutes. Afterwards, the suspension was centrifuged again for 15 minutes at 3500g. The same procedure was used following the second centrifugation as described for the first centrifugation step. For the 3^rd^ centrifugation, the time was shortened to 10 minutes. After the third and final centrifugation, the pellet was re-suspended in 0.5 ml of a 0.2 mg/ml solution of Bovine Serum Albumin (BSA) in DI water to block the remaining unreacted sites on the gold nanoparticles. The particles were then incubated on the orbital table for an additional 5 minutes. This was followed by the addition of 0.5 ml of DI water and 0.5% sodium azide.

### Detection of anti-Protein E antibody in vitro

To 100 µl of conjugated gold nanoparticles, 900 µl of PBS were added. Posteriorly, 10 µl of different concentrations of anti-Protein E antibody were added prior to measuring 90-degree light scattering using the oval plastic chamber and the handheld optical caustic sensor which used a Nokia 920 smartphone and Lumia software for imaging. Pictures of the oval chamber were taking using different times of exposure (0.1 sec or 1 sec) and by reducing overall brightness to the lowest level in the Lumia camera app. The images were analyzed measuring the raw intensity using ImageJ using the green or red channel, depending upon the level of saturation of the green channel, and by focusing in the zone of the sample chamber where scattering by the gold nanoparticles provides pixels with brightness above the typical PBS solution pixel values of 0-5 (out of a maximum range of 255).

### Dot Blot using conjugated Gold nanoparticles

Standard dot blot paper (Bio Rad Laboratories, Hercules, CA) was cut into rectangular or hexagonal shapes in order to fit into 12 well Corning Costar Tissue Culture Plates (Ted Pella Inc., Reading, CA). After punching a hole in the center of the paper with a sterile pin, 3 microliters of 1:100 PBS diluted human serum was placed over the area around the pin hole and allowed to dry for 5 minutes at 37 ˚C. Then, the paper was cooled to 4 ˚C and allowed to incubate for 15 minutes. Following incubation, an aliquot of 1 ml of a 3% skim milk in PBS blocking solution was added to the chamber and mixed for 15 minutes at room temperature. After incubating in the blocking solution for 30 minutes at 37 C, an aliquot of 500 microliters of a 1:10 diluted suspension of 60 nm Protein E conjugated gold nanoparticle was added to the paper. Incubation with the gold nanoparticle conjugate was conducted for 1 hour at 37 ˚C. After the final incubation, the paper was rinsed 3 times in PBS – Tween 20 buffer and allowed to dry.

### Testing of Human Serum Samples

All human serum samples were obtained voluntarily from Hemolab S.A. and the Lab of Arbovirology of the Universidad Autonoma de Yucatán collected for previous studies. Two 6 ml bottles of conjugated gold nanoparticle reagent GNP-60-E were created according to the protocol described above except that while one GNP reagent (GNP-60-E-3) was centrifuged 3 times to remove unbound Protein E, a second reagent (GNP-60-E-1) was centrifuged only once. Both reagents were stored for 16 hours at 4 ˚C then transported at room temperature for 19 hours, until refrigerated at 4 ˚C at the Universidad Autónoma de Yucatán (UADY). Both GNP-60-E reagents arrived un-aggregated at UADY based on both Nanodrop spectrometer (Bio Rad Laboratory, Hercules, CA) visible spectra taken at the Universidad Autónoma del Yucatán (UADY) before using the reagents and by visual observation of the gold nanoparticle suspension.

For each test, in a 1.5 ml Eppendorf tube, 900 µl of PBS were combined with 100 µl of the gold nanoparticle conjugate followed by the addition of 5 or 10 microliters of the human serum samples or a negative control of PBS. For the testing at 37 ˚C, GNP-60-E-3 was used and 10 microliters of undiluted human serum was added. Incubation was performed for 30 minutes at 37 ˚C without agitation. Afterwards, each test sample was placed in an oblate spheroid chamber and capped, followed by imaging with the smartphone sensor. For the first trial at 22 ˚C, GNP-60-E-3 was used and 5 microliters of undiluted human serum was added. Incubation was performed for 15 minutes at 22 ˚C without agitation. Afterwards, each test sample was placed in an oblate spheroid chamber and capped, followed by imaging. For the second trial at 22 ˚C, GNP-60-E-1 was used and 2 microliters of 1:10 diluted human serum was added. Incubation was performed for 15 minutes at 22 ˚C without agitation. Afterwards, each test sample was placed in an oblate spheroid chamber and capped, followed by imaging.

### Light Scattering Measurement Systems

For 90-degree laser light benchtop scattering measurements, an Ocean Optics Fiber Optic Spectrometer (Ocean Optics, Dunedin, FL) was used with a USB4000 spectrometer and SpectraSuite Software for control and data acquisition. A quartz cuvette of 1 cm path length with all four windows clear for measurement enables 90-degree placement of an InPhotonics 532 nm laser, with an Ocean Optics controller from the fiber optic collector, to measure counts of scattered light. Readings of 1 second integration time with 5 spectra averaging was used.

### 3D Printed Oblate Spheroid Chamber

Autodesk Fusion 360 software was used to generate the 3D oblate spheroid sample chamber used in the handheld optical caustic sensor. In order to place the sample chamber in the LED illumination chamber and easily remove it after imaging, a solid rectangular peg was added to the center of the base. The opening of the sample chamber was sized and designed with a small ridge in order to accommodate lids cut from 200 microliter PCR tubes. Fabrication of the sample chambers were done in batches of 12 by transmitting the drawing to a 3D printing company (iMaterialise, Leuven, Belgium). 3D printing is conducted using stereolithography with a transparent resin and the finished product is between water clear and translucent, in terms of visibility to the naked eye.

### Handheld Smartphone Optical Caustic Light Scattering Sensor

A previously described handheld smartphone-enabled optical caustic light scattering sensor was used to capture images of samples in an oblate spheroid chamber [1]. A 532 nm green Photodiode (Industrial Fiber Optics, Tempe, AZ) illuminated the sample chamber at a 90˚ angle from a Nokia Lumia 920 (Nokia, Espoo, Finland) smartphone camera lens. Images were collected at 0.5 second exposure using the Lumia camera app. At this exposure level, and with the intensity of light used, the green channel of the RGB images saturate. However, excess light due to scattering can be imaged in the center of the sample chamber by splitting the color channels using ImageJ (NIH, Bethesda, MD). ImageJ software was used to collect the intensity of the red, green, and blue channels for each image. Integrated density measurements from ImageJ was collected for a constant image area and used to quantify the amount of scattering observed for gold nanoparticles in water, conjugated gold nanoparticles with a serum sample, or a Phosphate Buffered Saline (PBS) control.

### Digital Color Optical Caustic Light Scattering Sensor

In lieu of a smartphone to collect images and in order to corroborate the gold nanoparticle scattering vs. size calibration with the optical caustic sample chamber, a Digital Color Sensor S9706 (Hamamatsu Photonics, Hamamatsu, JP) collected light from the oblate spheroid sample chamber at a 90˚ angle and a distance of 10 mm from the center of the chamber. Due to the proximity of the sensor to the sample chamber, a 3-D printed tubular mask with an opening of 7 mm in diameter was used to block the incident light. A 3-D printed chamber with interior angled walls was also fabricated and used to minimize light reflection. The digital color sensor was connected to an Arduino Uno computer and the manufacturer’s recommended algorithm was deployed to acquire data at different integration times. Data for the gold nanoparticles in water was acquired using between 0.1 - 30 seconds integration time. Since a lower intensity of LED light was used than with the smartphone system, data for 20, 40, 50, 60, 80, 100, and 200 nm was collected for the green channel of the sensor. The most accurate data to compare scattering using the digital color sensor in this system was the highest integration time of 30 seconds, which was then used to compare with the smartphone light scattering results.

## Results and Discussion

Before conjugation of Protein E to gold nanoparticles, the choice of gold nanoparticle size was deemed to be a key variable that needed to be determined. Based on the well understood properties of gold nanoparticles to light scattering via plasmon resonance [6] near the maximum for spheres of 520-540 nm, a calibration was conducted with a laser light scattering fiber optic spectrometer, the optical caustic sensor, and calculations based on Mie Theory [6]. Figure 1 illustrates two key points. First, that the laser light scattering system which is based on 90 degree scattering is sensitive to small changes in particles size at a narrow range of between 30-70 nm diameter particles. Mie Theory calculations illustrate that the total scattering for gold nanoparticles increases more gradually with size, since this calculation takes into account all angles of scattering. The Optical Caustic sample chamber is seen in Figure 1 to fall between the two groups of data, using both a camera and a digital color sensor to collect data. It was also found that the red channel of the smartphone RGB image was most useful based on the LED intensity and shutter speed setting since smartphone sensor saturation “bleeds” signal to the red and blue channels when there is an excess of green in the image. The digital color sensor corroborates that interpretation, as shown in Figure 1.

**Figure 1.**
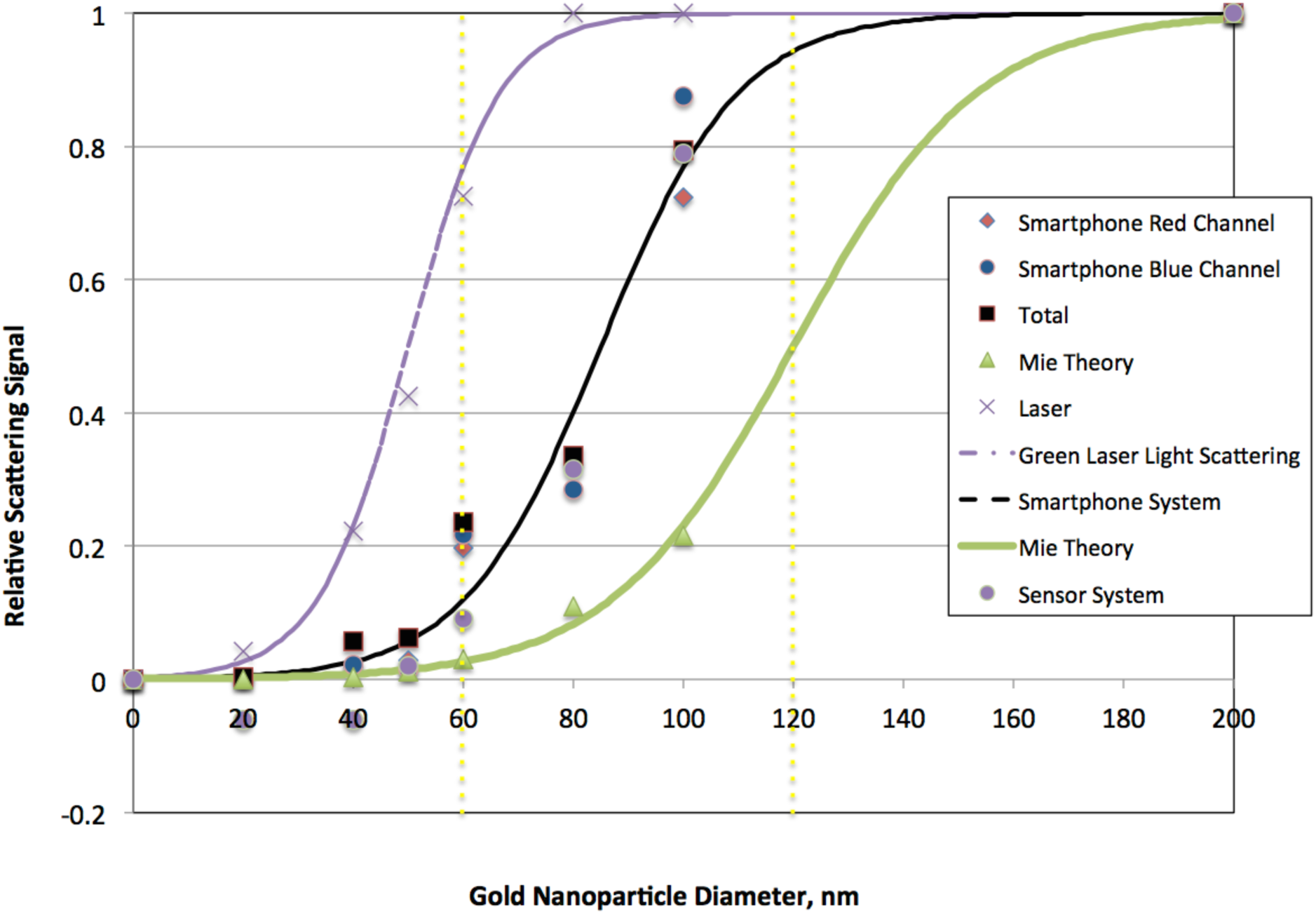
Comparison of gold nanoparticle scattering with green light data for a laser and fiber optic spectrometer, LED/optical caustic smartphone, and LED/optical caustic digital color sensor. Discrete calculations of the total scattering at 532 nm using Mie theory is also shown. Data and calculations are normalized to the maximum value for gold nanoparticles of 200 nm and fitted to a logistic curve. Horizontal dashed lines illustrate the signal difference for 60 and 120 nm diameter gold nanoparticles.

The overall interpretation of the calibration data is that using 60 nm gold particles is very practical for the intended application. The reason for this is that there is a dramatic (approximately 340%) increase in signal predicted when two 60 nm gold particles aggregate upon antibody-antigen binding, basically the change in relative signal from 60 nm to 120 nm gold particle size.

After establishing that 60 nm gold particles would be a useful size, it is also important to note that conjugated gold nanoparticles are sensitive optically to any surface change, including binding of antibodies or antigens to their surfaces and the additional binding of proteins to their surfaces or aggregation of gold nanoparticles. To establish the response of the system in terms of 90-degree light scattering near the plasmon resonant peak, different amounts of anti-Protein E antibody were added to a diluted solution of conjugated gold nanoparticles. The first parameter to determine was the zone for no scattering in the chamber, and in this case PBS without gold nanoparticles was used. Images taken with 1 sec of exposure and analyzed using the red channel since the green channel was already saturated with green light (Figure 2B) shows that when using PBS, the amount of light detected from the center of the sample chamber is very low since no scattering nor reflection is expected in this zone of the sample chamber. Other images using different exposure times with the green channel did not provide a sufficiently good baseline dark zone, so the most appropriate exposure time for imaging for the handheld optical caustic smartphone was determined to be a 1 sec exposure in the center zone of the images’ red channel.

**Figure 2.**
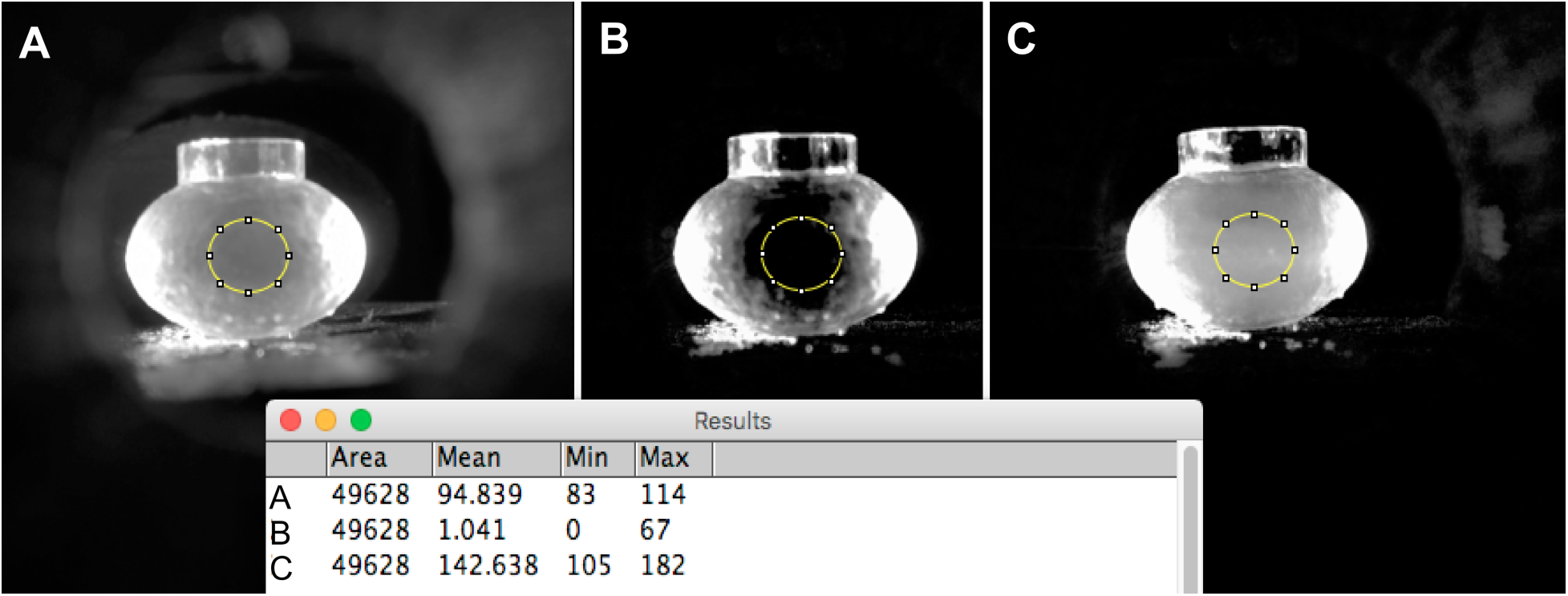
Light scattered using PBS or gold nanoparticles in the oval chamber. A) Light scattering intensity of PBS measured with the green channel of a picture taken with 0.1 sec of exposure time. B) Light scattering intensity of PBS measured with the red channel of a picture taken with 1 sec of exposure time. C) Light scattering intensity of 60 nm gold nanoparticles measured with the red channel of a picture taken with 1 sec of exposure time.

The table in the figure represent the gray values for the area inside the circle for each sample.

Next, images were taken for an aqueous suspension of 60 nm gold nanoparticles using the same imaging settings as for PBS. Figure 2C shows that the region given as a dark area when imaging PBS, now contains brightness due to gold nanoparticle light scattering. The integrated difference for these two measurements is on the order of 100x, which clearly provides a high range for detecting small changes in gold nanoparticle scattering. It is important to note that this signal is based on only ~1.8 billion gold nanoparticles in 0.7 ml of suspension, which suggests that detection to a sensitivity of 50 pg/ml of antibody or antigen should be possible since it could represent a 1% difference in nanoparticle scattering signal.

Finally, in order to quickly explore the range of signal change possible for a commercially available Protein E antigen with the handheld optical caustic sensor, the light scattering intensity for a gold nanoparticle suspension and gold nanoparticles containing Protein E in the presence of 10 µg/ml of anti-Protein E antibody in PBS. Table 1 gives the gray value differences for the controls and the antibody test.

**Table 1.**
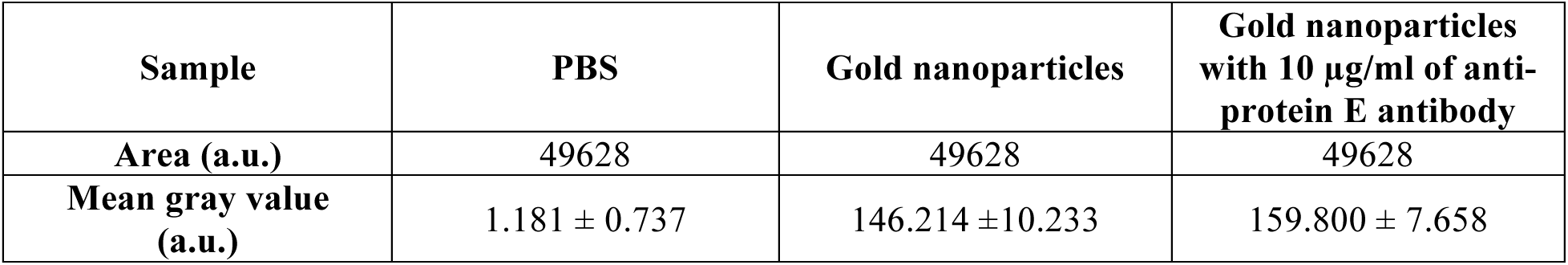
**Average gray values as a light scattering measurement for samples.** Mean gray values for samples of PBS, gold nanoparticles, and gold nanoparticles with 10 µg/ml of anti-protein E antibody, in the oval chamber.

The initial challenge lay in interpreting patient results without an extensive study of the antigen protein structure nor its immunogenic properties when presented on gold nanoparticle surfaces. This makes discerning the performance of the tests with respect to patient Dengue status characterization challenging. The challenge is somewhat reduced by using a dot blot test with the same patient samples and gold conjugate reagents as an orthogonal test. Table 2 provides information on the 10 patient serum samples donated by the Arbovirus laboratory of UADY for this study. These samples represent a spectrum of patients including Dengue negative, Dengue serotype 2 positive with a first time (primary) infection and 2^nd^ time (secondary) infection, patients convalescing from a Dengue serotype 2 infection, and patients with Dengue serotypes 1 and 3, respectively. Patients with early stage and primary dengue infection should present IgM against dengue antigens after a few days of infection and the IgM levels should decrease after 90 days [3–5, 8–11]. Patients with a secondary infection should mostly present IgG antibodies [5 8–11]. The antigen Protein E used as the reagent conjugated to the gold nanoparticles is a recombinant protein with selectivity towards Dengue Serotype 2 antibodies and is described by the manufacturer as being most sensitive to IgM levels. However, it is known that there can be cross-reactivity to other dengue serotype antibodies as well as from patients who have experienced other arbovirus infections from the group known as flaviviruses such as West Nile, St. Louis Encephalitis, Zika Virus, and Chikungunya [4]. These considerations suggest that there should be some variation in test results, but the variations may be difficult to discern without a more comprehensive series of human serum controls.

**Table 2.** **Description of human serum samples.** Human serum samples used in the feasibility study with descriptive information on Dengue status.

| Patient | Dengue Status | Description of Sample | Primary or Secondary Infection | Stage of Disease |
| --- | --- | --- | --- | --- |
| 1 | Negative | Asymptomatic, healthy individual, PCR-IgM-IgG negative for Dengue Virus | N.A. | N.A. |
| 2 | Negative | Asymptomatic, healthy individual, PCR-IgM-IgG negative for Dengue Virus | N.A. | N.A. |
| 3* | DENV-2 | PCR positive for DENV-2, IgM positive for DENV-2, IgG negative for DENV-2 | Primary | Initial stage |
| 4* | DENV-2 | PCR positive for DENV-2, IgM positive for DENV-2, IgG negative for DENV-2 | Primary | Initial stage |
| 5 | DENV-2 | Seropositive for DENV-2, IgM negative for DENV-2, IgG elevated for DENV-2, day 4 | Secondary | Initial stage |
| 6 | DENV-2 | Seropositive for DENV-2, IgM negative for DENV-2, IgG elevated for DENV-2, day 4 | Secondary | Initial stage |
| 7 | DENV-3 | PCR positive for DENV-3 | Unknown | Initial stage |
| 8 | DENV-1 | PCR positive for DENV-1 | Unknown | Initial stage |
| 9** | DENV-2 | Volunteer donated sample, seropositive for DENV-2, IgM positive for DENV-2 | Unknown | Convalescing > 120 days |
| 10** | DENV-2 | Volunteer donated sample, seropositive for DENV-2, IgM positive for DENV-2 | Unknown | Convalescing > 120 days |

\*Patients positive for dengue with negative IgG test.
\*\*Patients volunteered for testing were positive with DENV-2 during the epidemic in 2015 and donated samples 4 months after initial diagnoses.

Before commencing patient testing, the 60 nm gold conjugates were inspected and the UV/VIS spectra were recorded for the two batches of particle conjugates (Figure 3). Both visual and spectral information verified that the gold conjugates were stable and ready to use at UADY.

**Figure 3.**
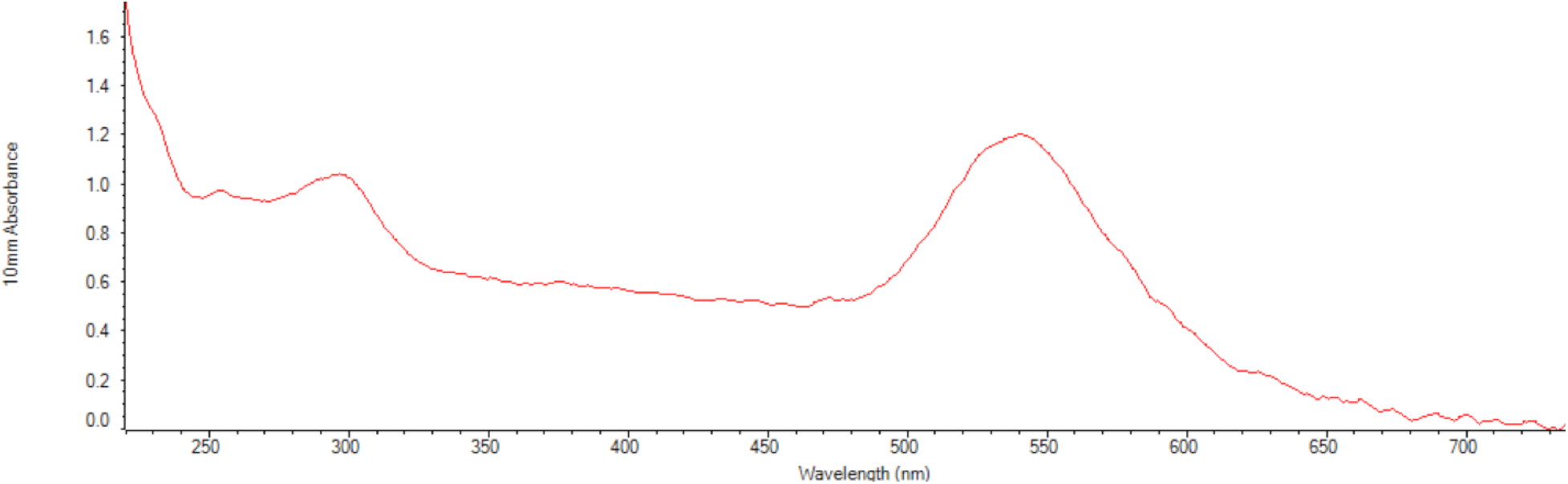
Nanodrop spectrometer UV/VIS spectra. The absorbance in the UV is due to the BSA blocking solution. The absorbance maxima at a wavelength of 540 nm is consistent with the expected plasmon resonant peak for conjugated gold nanoparticles of 60 nm diameter.

Table 3 summarizes the expected dot blot score based on the protein E gold conjugate reagent (GNP-60-E-3) used, assuming some level of cross-reactivity with other Dengue serotypes and a 50% probability of measuring antibodies reactive to Protein E after 120 days of convalescence. The individual blot samples and the scores of the dot blot are shown in Figure 4 and in Table 3.

**Table 3.** **Comparison of results of the human serum samples between diagnostic methods.** Comparison of Dot Blot with Expected Result due to Dengue Status reported by the clinical laboratory. The Protein E used while specific for DENV-2, has sensitivity for IgM detection and can have cross-reactivity with other serotypes.

| Patient | Expected Dengue Result Based on Dengue Serotype and Some Cross-Reactivity (assuming high sensitivity to IgM) | Dot Blot Score ( + or -) | Agreement |
| --- | --- | --- | --- |
| 1 | - | + | NO |
| 2 | - | + | NO |
| 3 | + | + | YES |
| 4 | + | + | YES |
| 5 | - | - | YES |
| 6 | - | - | YES |
| 7 | - | - | YES |
| 8 | + | + | YES |
| 9 | + | + | YES |
| 10 | - | - | YES |

**Figure 4.**
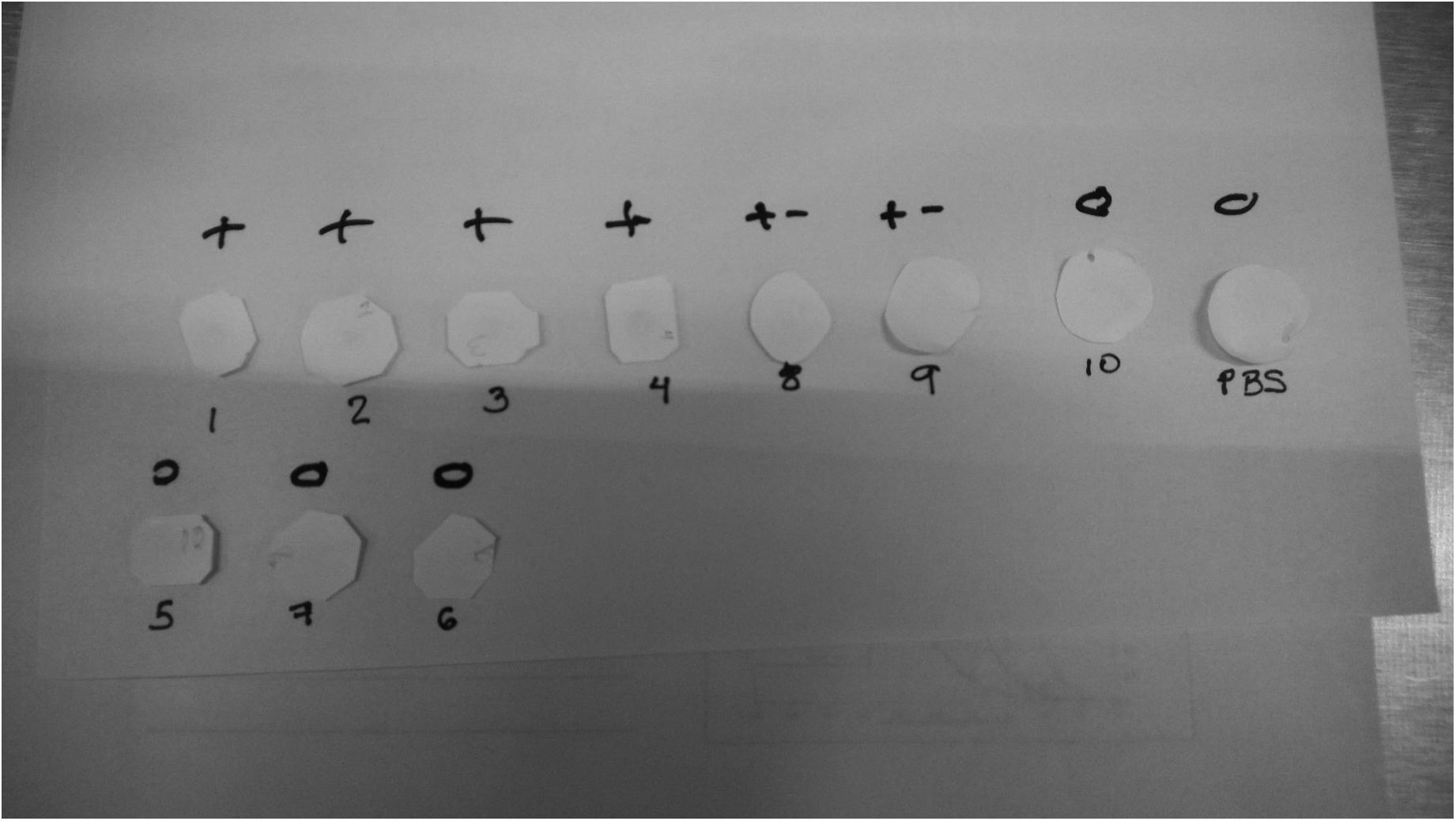
Image showing visual scoring of dot blots. Each blot was done in separate paper and reaction chambers in order to limit solution volume for gold conjugation and prevent cross contamination among the human serum samples and PBS negative control.

Overall, the agreement is very good. However, the two dengue negative patients appear to give positive dot blot results. It is not clear why this is the case, but perhaps a current or prior flavivirus infection is being detected by the gold conjugate reagent. Additionally, the sample with PBS added and no human serum was correctly identified as a negative control in the dot blot experiment. It is also important to note that a dot blot test (not shown) with a monoclonal antibody that reacts with flavivirus group specific antigens (4G2, KPL) was found to be negative. The negative result suggests that epitopes of the protein envelope needed for the monoclonal antibody to bind to the GNP-60-E-3 were blocked due to the immobilization of Protein E to the gold nanoparticles. This is a potentially useful observation for further gold conjugate development, namely if a methodology for Protein E binding to gold based on covalent attachment rather than his-tag is used.

A summary of the three sets of tests with human serum samples and an overall score is given in Table 4 while a quantitative comparison of the dot blot is shown in Figure 5. The overall score given in Table 3 is based on assessing the differences among the three tests using gold nanoparticles made with either 3 or 1 centrifugation step(s) and at different times and temperatures. Based on the results in Table 3, it appears that the results with GNP-60-E-1 at 22 C for 15 minutes seem to be most closely related to the characterization based on the dengue patient status information.

**Table 4.** Qualitative summary of Optical Caustic Scattering Smartphone Sensor data.

| **Patient** | **GNP-60-E-3**<br>**37 C for 30 min** | **GNP-60-E-3**<br>**22 C for 15 min Trial 1** | **GNP-60-E-1**<br>**22C for 15 min Trial 2** | **Overall**<br>**Score (+ or -)** |
| --- | --- | --- | --- | --- |
| **1** | weak positive | *positive* | negative | - |
| **2** | positive | *negative* | positive | + |
| **3** | positive | *positive* | positive | + |
| **4** | positive | *positive* | weak positive | + |
| **5** | positive | *positive* | positive | + |
| **6** | positive | *negative* | positive | + |
| **7** | negative | *weak positive* | negative | - |
| **8** | positive | *positive* | negative | - |
| **9** | positive | *weak positive* | positive | + |
| **10** | very weak positive | *positive* | positive | + |
| **PBS Control** | negative | *negative* | negative | **NEGATIVE**<br>**CONTROL** |
| **PBS Control** | negative | *negative* | negative | **NEGATIVE**<br>**CONTROL** |

**Figure 5.**
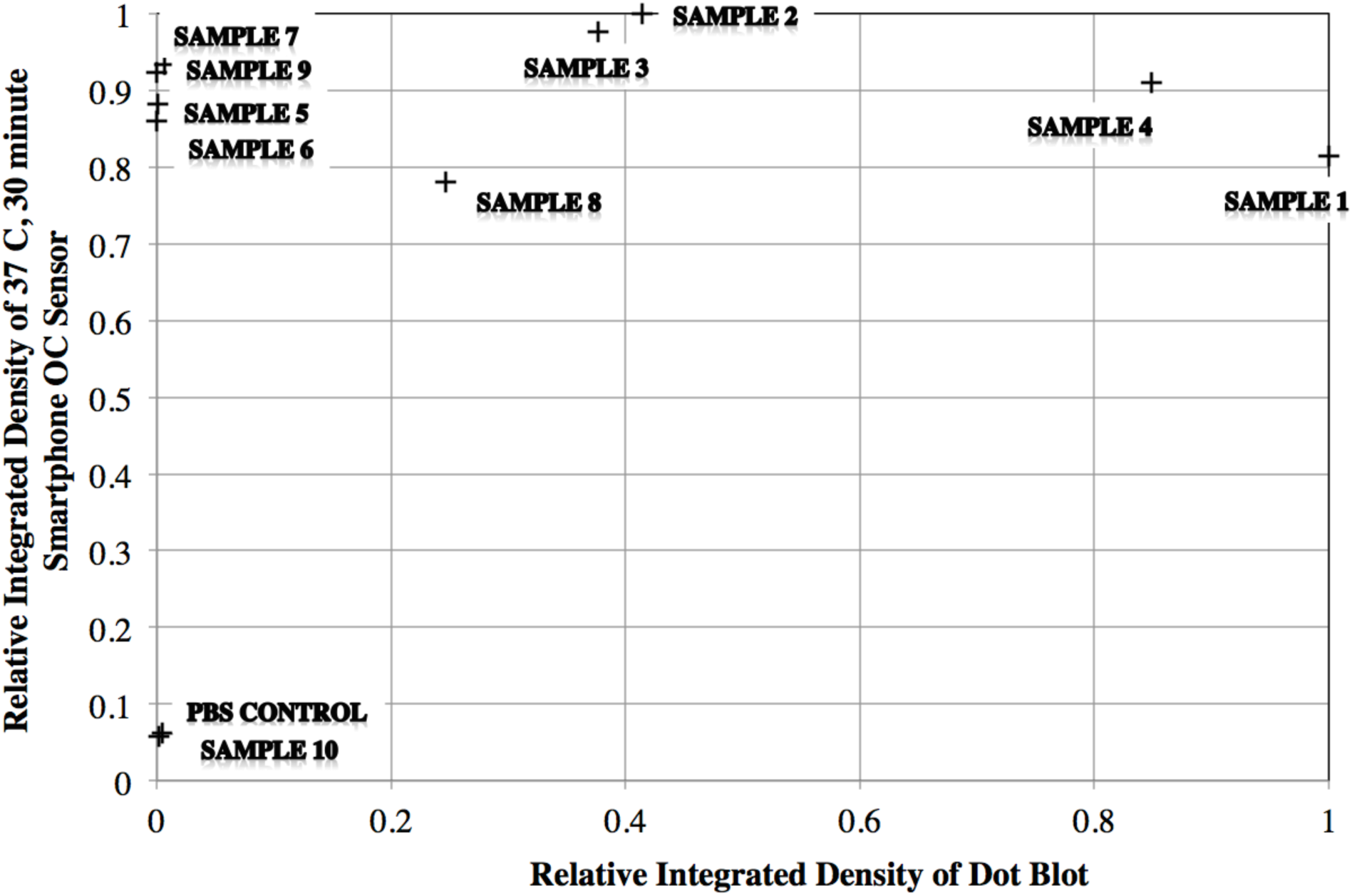
Comparison of Dot Blot and Optical Caustic Smartphone Sensor. Data were recorded at the same sample incubation temperature of 37 C.

In Figure 5, there are two overall features to note. First it appears that the smartphone signal saturates at low “signals” from the dot blot experiments. This seems reasonable given the high signal change described in Figure 1 and 2 for aggregates consisting of 60 nm gold particles. The second feature is that the negative control and patient sample 10 are both low for the dot blot and the optical caustic sensor, but there is disagreement between patient samples 5 and 6 between the two detection methods. This may be due to the higher sensitivity of the optical caustic sensor as compared to the dot blot which used gold nanoparticles at a higher level of dilution. Dot blots are also considered to be lower in sensitivity due to diffusion and orientation effects when binding to a fixed surface.

Another observation is that patient sample 7 yielded a very weak signal and patient sample 10 is negative based on the dot blot tests, yet the optical caustic sensor data show strong signals. This again may be due to higher sensitivity of the optical caustic system and cross reactivity of the Dengue Serotype 2 recombinant Protein E with other Dengue serotypes. As noted in the results for GNP-60-E-1 at 22 C and 15 minutes seen in Table 4, it appears that this cross-reactivity may be diminished by using a higher dilution of human serum, lower temperature, and allowing some unbound antigen in the gold nanoparticle conjugation suspension.

## Conclusions

A feasibility study of a hybrid gold nanoparticle with an optical caustic sensor system indicates that the technology is sensitive to the detection of serum antibodies against dengue fever. The use of similar size gold nanoparticles (60 nm) to what is often used in paper lateral flow rapid tests is justified based on calibration data and human patient sample testing. The gold nanoparticle conjugates are stable to transportation and temperature fluctuations and appear to be un-aggregated and active after several days of conjugation. Also, all negative controls for the dot blot and optical caustic sensors were correctly measured as negative.

Dot blot tests were found to be useful to verify the reactivity of the gold nanoparticle conjugates to patient serum antibodies and may also help identify epitope presentation differences when applying monoclonal antibodies to the dot blot paper directly. It appears that human serum should be diluted and that testing can be done at around 22 ˚C within 15 minutes using the hybrid gold nanoparticle conjugate reagent and optical caustic sensor. A more thorough investigation of well characterized human serum samples is still needed in order to optimize the test conditions and buffer components and generate positive predictive and negative predictive values that reflect the capabilities of the diagnostic platform.

## Acknowledgements

The authors would like to thank the universities involved in the project, namely: Instituto Universitario Puebla S.C. and Instituto de Consultoria Universitaria Santin SC, for their participation. We also acknowledge Hemolab S. A. de C. V. and Lab of Arbovirology of the Universidad Autonoma de Yucatan for the human patient samples and testing to provide information on their Dengue status, which were obtained before the research study began.

## References

1. García A. A., Luis Nuñez, Angel Lastra, and Vladimiro Mujica, Application of Newton’s Zero Order Caustic for Analysis and Measurement: Part-III Light Scattering, International Research Journal of Pure and Applied Chemistry, ISSN: 2231-3443, Vol.: 4, Issue: 1 (January-March) Page 144–158. (2013)

2. Lipowicz, Michelle, García, A. A., Handheld Device Adapted to Smartphone Cameras for the Measurement of Sodium Ion Concentrations at Saliva-Relevant Levels via Fluorescence, Bioengineering 2015, 2(2), 122–138.

3. Rockstroh A, Barzon L, Pacenti M, Palù G, Niedrig M, et al. (2015) Recombinant Envelope-Proteins with Mutations in the Conserved Fusion Loop Allow Specific Serological Diagnosis of Dengue-Infections. PLoS Negl Trop Dis 9(11): e0004218. doi: 10.1371/journal.pntd.0004218

4. Cuzzubbo AJ, Endy TP, Nisalak A, et al. Use of Recombinant Envelope Proteins for Serological Diagnosis of Dengue Virus Infection in an Immunochromatographic Assay. Clinical and Diagnostic Laboratory Immunology. 2001;8(6):1150–1155. doi:10.1128/CDLI.8.6.1150-1155.2001.

5. Lin H-E, Tsai W-Y, Liu I-J, Li P-C, Liao M-Y, et al. (2012) Analysis of Epitopes on Dengue Virus Envelope Protein Recognized by Monoclonal Antibodies and Polyclonal Human Sera by a High Throughput Assay. PLoS Negl Trop Dis 6(1): e1447. doi: 10.1371/journal.pntd.0001447

6. Fan X, Zheng, W, Singh, D, Light scattering and surface plasmons on small spherical particles, Light: Science & Applications (2014) 3, e179; doi:10.1038/lsa.2014.60

7. Katzelnick L.C., Fonville J.M., Gromowski G.D., Bustos-Arriaga J., Green A., et al (2015). Dengue viruses cluster antigenically but not as discrete serotypes. 1338 18 SEPTEMBER 2015 • VOL 349 ISSUE 6254 sciencemag.org SCIENCE

8. M.S. Mustafa, V. Rasotgi, S. Jain, V. Gupta (2015). Discovery of fifth serotype of dengue virus (DENV-5): A new public health dilemma in dengue control. Medical Journal Armed Forces India Volume 71, Issue 1, January 2015, Pages 67–70

9. da Silva Voorham JM (2014) A possible fifth dengue virus serotype. Ned Tijdschr Geneeskd. 2014;158:A7946.

10. Anne Tuiskunen Bäck, and Ake Lundkvist (2013). Dengue viruses-an overview. Infection Ecology and Epidemiology 2013, 3: 19839 - http://dx.doi.org/10.3402/iee.v3i0.19839

11. World Health Organization (2009). Dengue: guidelines for diagnosis, treatment, prevention and control -- New edition. Chapter 4 Laboratory diagnosis and diagnostic tests. ISBN 978 92 4 154787 1

